# A novel approach to the empirical characterization of learning in biological systems

**DOI:** 10.1101/2021.01.10.426118

**Authors:** Yarden Cohen, Predrag Cvitanović, Sara A. Solla

## Abstract

Learning to execute precise, yet complex, motor actions through practice is a trait shared by most organisms. Here we develop a novel experimental approach for the comprehensive investigation and characterization of the learning dynamics of practiced motion. Following the dynamical systems framework, we consider a high-dimensional behavioral space in which a trial-by-trial sequence of motor outputs defines a trajectory that converges to a fixed point - the desired motor output. In this scenario, details of the internal dynamics and the trial-by-trial learning mechanism cannot be disentangled from behavioral noise for nonlinear systems or even well estimated for linear systems with many parameters. To overcome this problem, we introduce a novel approach: the sporadic application of systematic target perturbations that span the behavioral space and allow us to estimate the linearized dynamics in the vicinity of the fixed point. The steady-state Lyapunov equation then allows us to identify the noise covariance. We illustrate the method by analyzing sequence-generating neural networks with either intrinsic or extrinsic noise, at time resolutions that span from spike timing to spiking rates. We demonstrate the utility of our approach in experimentally plausible and realizable settings and show that this method can fully characterize the linearized between-trials learning dynamics as well as extract meaningful internal properties of the unknown mechanism that drives the motor output within each trial. We then illustrate how the approach can be extended to nonlinear learning dynamics through a flexible choice of the basis and magnitude of perturbations.

**Significance statement:** Movement control ties brain activity to measurable external actions in real time, providing a useful tool for both neuroscientists interested in the emergence of stable behavior and biomedical engineers interested in the design of neural prosthesis and brain-machine interfaces. We approach the question of motor skill learning by introducing artificial errors through a novel perturbative scheme amenable to analytic examination in the linearized regime close to the desired behavior. Numerical simulations then demonstrate how to probe the learning dynamics in both linear and nonlinear systems. These findings stress the usefulness of analyzing responses to deliberately induced errors and the importance of properly designing such perturbation experiments. Our approach provides a novel generic tool for monitoring the acquisition of motor skills.

## Introduction

Many daily actions, such as visually guided reaches or handwriting, are forms of well-practiced motion. The execution of such skilled behavior is often described as optimized motor control – the creation of a desired movement trajectory, called target motion, by minimizing the deviation of ongoing behavior from that expected path (1–3). The neural circuits underlying both motor control (4), and learning and maintaining skilled behavior (5) are still poorly understood. In behavioral studies, the learning algorithms implemented by these circuits can be investigated by changing the target motion and forcing adaptation (6,7), re-optimization (8), or re-learning if the target was acquired in the past. These psychophysical experiments revealed that human subjects generalize the response to target change across the space of motor outputs (9), suggesting that the brain holds an internal model of the body’s physical plant (10) and can use this representation in behavior adaptation (11–13). The state-space theory describes these internal models using a set of motor primitives spanning all possible behavior outputs. This framework relates the adaptation to the target motion perturbation in a previous trial (14,15) and its generalization to new target motions (16).

The linear dynamical systems (LDS) approach used in (7,14–16) extends the state-space representation of a single motion to a sequence of motions by assuming linear relations between consecutive motor executions. The linearity assumption offers a powerful mathematical framework for comparing motor learning algorithms by fitting the trial-by-trial motor errors observed in experiments (17) and extracting the internal noise in the premotor drive. However, fitting learning behavior with LDS is a non-convex optimization problem that can be biased in important scenarios. Specifically, many experiments elicit within-trial response to perturbations (18,19), non-linear responses to large perturbations, or use frequent stochastic perturbations that can mask the internal noise sources. Additionally, repeating activity sequences have been observed in the CNS without direct connection to a known physical output or from an uncertain stage in the overall function (20,21). Neural network models producing these reliable activity patterns may have internal degrees of freedom and dynamics that are strikingly different from the behavioral observations (e.g. (22) for feed-forward and (23) for near chaos recurrent networks). In such cases, as well as in complex motor learning studies, it may be impossible to choose a small set of degrees of freedom and reduce the motor primitives state space used in LDS – an important step in fitting these models whose number of parameters is quadratic in the number of degrees of freedom.

Here we propose to augment the target motion perturbation approach to provide an experimental framework for exhaustive characterization of the learning dynamics around practiced motion trajectories. Specifically, instead of using frequent and large perturbations, we offer to use responses to sparse small perturbations that span a subset of generative primitives. Instead of investigating learning algorithms along all steps between naïve and skilled performance, we offer to fully characterize the linear approximation to the learning dynamics involved in maintenance of already-practiced behavior. Using rate-based and spiking neural networks and a gradient-descent learning algorithm as black-box experimental subjects we exemplify, in agreement with (14), that our approach captures the linear learning dynamics with significant noise of multiple sources and with an experimentally feasible number or trials. We demonstrate our method’s ability to investigate the internal properties of these toy models. We then provide examples showing how our approach can circumvent the limitations of LDS and be used to study nonlinearities by augmenting perturbation magnitudes and by choosing different perturbation types.

### The asymptotic characterization of a dynamical system

We consider a behavioral task that implies the control of a system (such as a limb) whose state at any given time point t is characterized by d degrees of freedom. The task takes place during a time interval of duration T. We follow the evolution of the system during the execution of the task at a time resolution δ t, and include the initial condition at t = 0. An observation of an instantiation of the task corresponds to a trajectory of length (1 + K) through the d-dimensional state space of the system, with K = T / δ t. Each instantiation of the task is represented by a point X in a space of dimension D = d (1 + K) In this space of all possible instantiations of the task, there is a target X^*^ that represents the desired task. Iterative learning is represented as a sequence of trials {X_n_} ∈ ℝ^D^; successful learning corresponds to a convergent sequence 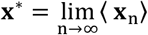. Here, the average ⟨…⟩ is over all possible initial conditions and realizations of the noise for the learning dynamics.

After sufficient training, the trial-by-trial task realizations are asymptotically close to the target **x**^*^. To characterize this asymptotic regime, we adopt the Linear Dynamical System (LDS) framework; this linearized approach has been applied to the modeling of sensorimotor learning (17). Stable convergence to the target implies that **x**^*^ is an attractive fixed point of the learning dynamics. The gradual convergence of the sequence of trials {**x**_n_} to the target **x**^*^ suggests the introduction of a new representation **y**_n_ = **x**_n_ − **x**^*^ that describes task realizations as deviations from the desired goal. The trial after trial dynamics of task acquisition near the fixed point **x**^*^ can then be linearized to obtain

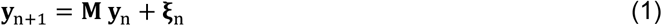

Here **ξ** _n_ is the noise, assumed to be an independent and identically distributed Gaussian D-dimensional random variable with zero mean and D × D covariance matrix **Δ, ξ**_n_∼ 𝒩 (0, **Δ**). The D × D matrix **M** implements the linearized dynamic. This matrix **M** could in general depend on the iteration number n, on the current state **x**_n_, or even exhibit a history dependence on a sequence of p preceding states **M**_n_(n, **x**_n_, **x**_n−1_, ..., **x**_n−p_), but here we assume that for large trial number n and sufficiently close to the fixed point, the matrix **M** is constant.

### The interplay between learning and noise perturbations

After sufficient learning to be within a neighborhood of the target, continued learning is likely to be required to maintain behavioral accuracy and prevent increased variability due to ever present motor noise, and to compensate for drift due to temperature or load fluctuations at intermediate time scales or to growth and aging over long time scales. Even at short time scales dominated by noise, stable behavior arises from the interplay between convergent learning and noise perturbations.

The variability of asymptotic learning is characterized through **y**_n_ = **x**_n_ − **x**^*^, with dynamics **y**_n+1_ = **M y**_n_ + **ξ**_n_ The noise averaged mean ⟨ **y**_n_ ⟩ satisfies the deterministic equation ⟨**y**_n+1_⟩ = **M** (**y**_n_), while variability is characterized through the second order statistics 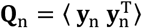. In the steady state around the fixed point ⟨ **y**_n_ ⟩ = 0, the covariance matrix ***Q*** satisfies the Lyapunov equation (see Methods)

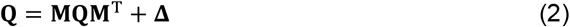

In this equation, the matrix **Q** is obtained by averaging the outer product 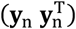 over many trials in steady state, a procedure equivalent to that of averaging over noise realizations. However, knowing **Q** does not suffice to untangle the contribution of the asymptotic dynamics **M** from that of the noise **Δ**.

In the study of dynamical systems, the linearized dynamics **M** are usually known (24,25) and the steady-state Lyapunov equation (Eq. 2) can be used to extract the noise covariance **Δ**. In the study of learning in artificial systems, known as machine learning, the learning algorithm is known and its linearized form **M** can be calculated. In contrast, in the case of learning in biological systems, the dynamics of learning are unknown and independent characterizations of the associated noise are unavailable. Here we show how a sequence of errors {**y**_n_} ∈ ℝ ^D^ can be used to estimate the components of the linear dynamics. The approach exploits the interplay between contracting dynamics and expanding noise, thus avoiding the ambiguities likely to arise in the noiseless case when only the asymptotic value **x**^*^ is known.

### A systematic application of external perturbations untangles deterministic dynamics from noise

The behavior sequence {**y**_n_} can be used for accurate estimation of the linear dynamics **M** (9,26) if a relatively small basis of the motor ‘state-space’ is known (14,16,17) and if the learning indeed follows a linear update algorithm. With no prior knowledge to allow for a reduction of the dimensionality D of the space in which the learning dynamics occurs and without prior knowledge of the learning algorithm, we can implement an experimental approach: to perturb the desired target for one iteration: **x**^*^ → **x**^*^ + ∈ **v** and measure the behavior, **x′**, in the next iteration. Here **v** is a unit vector of arbitrary orientation in the D-dimensional space, and E is small enough so that a linearized description of the learning dynamics with a constant dynamic matrix **M** is not affected.

After one perturbed trial, we allow for several unperturbed trials, during which the system is allowed to converge to **x**^*^ Averaging over several repetitions of this perturbation procedure using a fixed perturbation ε **v** leads to (see Methods),

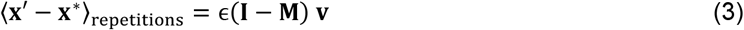

This scheme allows us to compute the effect of **M** on ***v***; repeating the procedure for a spanning basis of vectors {**v**} leads to the linearized dynamics matrix **M** The Lyapunov equation (2) can then be used to obtain the noise covariance matrix **Δ**, thus completing the characterization of the linearized dynamics. We note that this framework can be applied to any subset of perturbation vectors {**v**}, and as such allows the characterization of the linearized dynamics in specific subspaces as necessitated by the experiment.

### Learning dynamics of neural network models

We now apply the proposed method to simulated neural network models based on either rate or integrate-and-fire neurons. We also discuss the application of the method to nonlinear scenarios.

A leading hypothesis in motor control is that a high-dimensional internal dynamics of a network of neurons controls a relatively low-dimensional output. As the simplest example, consider a single readout neuron that outputs a scalar linear combination of the activity of all neurons in a network, 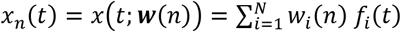. Here 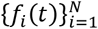 are the outputs of the N neurons in the network at time *t*, and 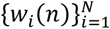 are the weights connecting the neurons in the network to the readout neuron on the n-th trial. The output trace during this trial is ***x***_*n*_(*t*), a vector in ℝ ^*K*+1^ parameterized by the discrete time index *t* ∈ [0 ..*K*] that spans the duration of a trial.

As in (14), we assume that the learning process converges to a desired target 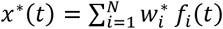 through gradient descent on the squared error with a constant learning rate *η*. This leads to the linear dynamic matrix (see Methods)

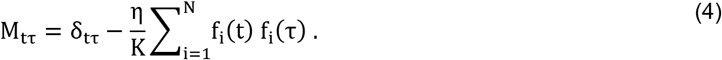

Note that the equation above holds regardless of the type of internal network dynamics and the nature of the outputs 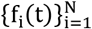. The network dynamics could be based on either rates or spikes; if we consider it to be deterministic, the time evolution of the neural network will depend only on its initial conditions. Note also that the learning dynamic matrix **M**_tτ_ is a (K + 1) x (K + 1) real and symmetric matrix; all its eigenvalues will be real, and the convergence of the learning process to the target **x**^*^(t) will involve exponential decay with no rotations. To model a realistic learning scenario, we must include noise in both the initial conditions and the network dynamics, as exemplified in the cases below.

### Example 1: a recurrent network of firing rate neurons

A recurrent network of N neurons characterized by their instantaneous firing rates 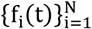 is often used as a substrate for sequence learning. It can produce a large variety of smooth outputs by adjusting the weights of the connections to a readout output neuron (23), given a large enough number N of neurons in the network and a sufficiently strong connectivity among them. The neural network used for these numerical experiments is described in Methods. The weights 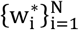 of the linear combination of neural firing rates that achieve the desired scalar output, 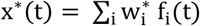, follow from gradient descent on the squared error. In this scenario, and in the absence of noise, the convergence of the learning algorithm to the desired solution is guaranteed by the stability of the learning dynamic matrix in Eq. 4. The questions to be investigated are the effect of network noise and of variability in network initial conditions on the matrix **M**, and the effect of noise on the convergence of the learning algorithm and on the ability of the perturbation method of Eq. 3 to accurately estimate the matrix **M**.

### The effect of initial condition on the linearized learning matrix M

The stability of the linearized learning dynamics (Eq. 1) across trials is an underlying assumption in our framework. The matrix **M**, calculated in Eq. 4 for the particular scenario now being considered, is trivially stable if we reset the network to fixed initial conditions in each trial and simulate noiseless dynamics (Eq. 6, Methods). But such resetting decouples one learning trial from the next. In the following sections we simulate successive trials of duration T and inter-trial intervals of the same duration using continuous noisy dynamics of the recurrent network (Methods).

The question that arises is that of the proper initial condition at the onset of successive trials. One possible choice is to use the current value of the neurons’ state variables 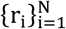 as the initial condition (here, the firing rate is a sigmoidal function of these variables, *f*_*i*_ = tanh (r_i_), see methods). This choice effectively introduces randomness in the initial conditions across trials, but it overlooks the existence of a neural ’go-signal’ that marks the beginning of each trial. We therefore choose to signal trial onset by the addition of a constant activation r_0_ to all 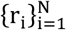 at the beginning of each trial (see Methods). This pulse of increased activation is the same for all neurons and across all trials, and its application at trial onset primes each trial snippet with a degree of commonality in initial conditions (see Fig. S 1 and SI section “Initial conditions, ‘go’ signal, and steady state dynamics of the rate-based network simulations”).

With the ‘go-signal’ we ensure reliability of the network’s dynamics and, as a result of Eq. 4, the stability of the matrix **M** across repeated trials, demonstrated through the stability of its eigenvalues and eigenvectors (Fig. S 2). In the following sections we compare the matrix **M**, estimated by the perturbation method, to its expected value from initializing a noiseless network at r_i_ = r_0_, ∀ i ∈ {1,... *N*}.

### The effect of noise

Various sources can introduce noise into a dynamical system. For our purposes of extracting the linearized learning dynamics we simulate noisy processes by adding independent zero-mean Gaussian noise with standard deviation σ to the network’s equation of motion (Eq. 6, see methods). This noise creates two effects that can limit using our perturbation method (Eq. 3). First, the added noise prevents the learning dynamics from converging, on average, to the desired target motion – a failure of the learning process itself that violates the condition 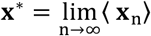 (Fig. S 3C, the average bias from the desired target, ∥ ⟨ **x**_n_ − **x**^*^⟩∥, was 1%-25% of the target’s Euclidean norm, ∥ **x**^*^ ∥, for σ = 0 → 0 01). Second, the noise increases the trial-to-trial output fluctuations around the fixed point (Fig. 1A, Fig. S 3D, the mean distance, ⟨ ∥ X_n_ − ⟨ X ⟩ ∥ ⟩, was 5-35% of the target’s Euclidean norm, ∥ **x**^*^ ∥, for σ = 0 → 0 01. The lowest value, 5%, resulted from the difference in initial conditions across consecutive trials). We next tested how these output bias and variability impact the success of averaging responses to perturbations (Eq. 3) in extracting the noiseless, fixed-initial conditions, dynamics matrix (Eq. 4).

**Fig 1.**
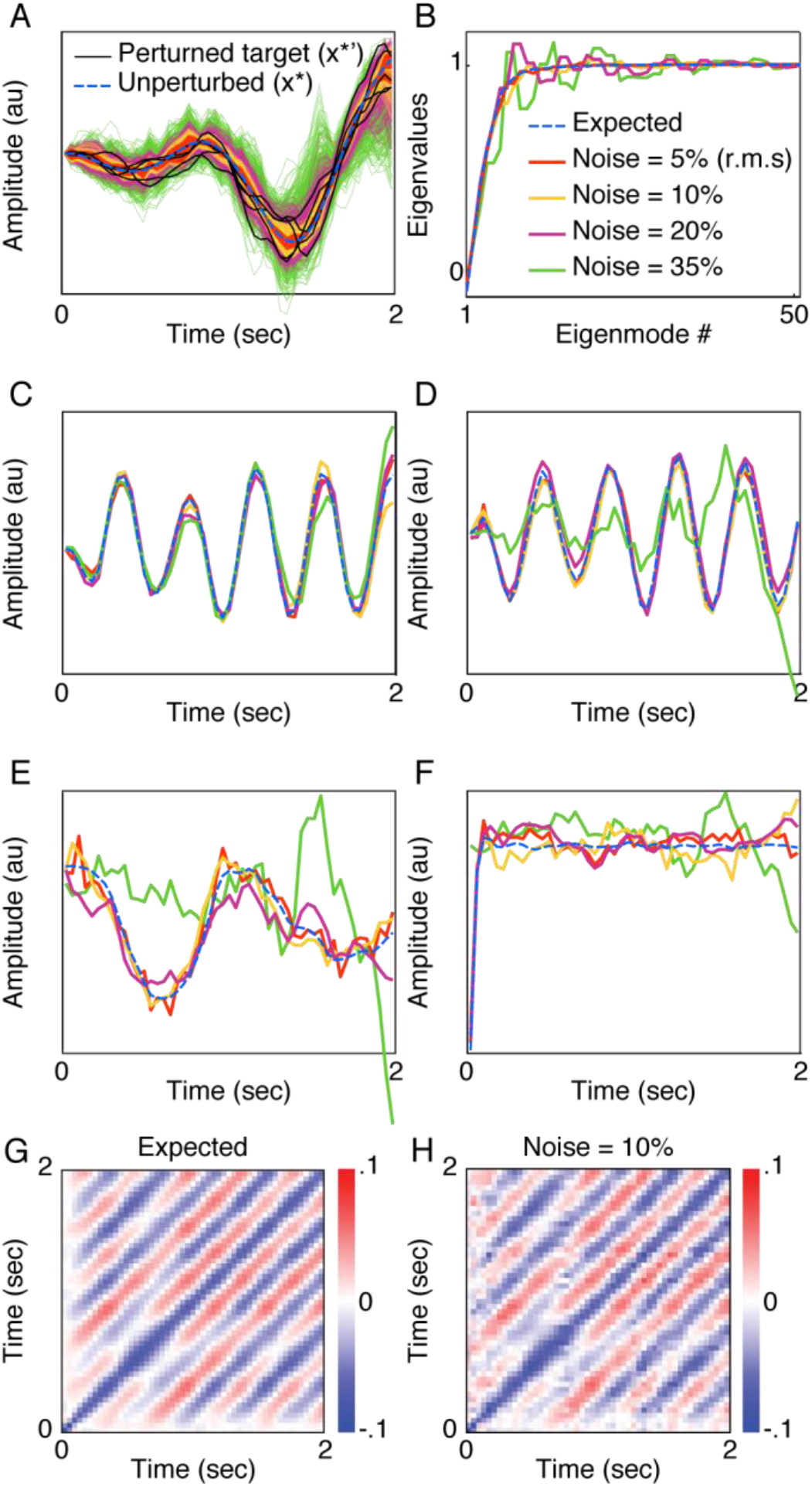
Perturbation method applied to a noisy network of firing rate neurons. **A**. The dashed blue line illustrates the desired target as a 2s sequence of values for the scalar output **x**^*^(t). Colored lines show consecutive noisy trials after the learning converged and before the perturbations started. Different colors stand for different noise values matching the legend in panel B. The black lines illustrate target perturbations, **x**^*^**′**(t). **B**. The spectrum of the linear dynamics matrix, **M**, expected in a noiseless system (dashed blue) and estimated by the perturbation method (colored). The different colors mark variance levels of the Gaussian noise added to Eq. 6 and resulting in trial-to-trial fluctuations measured by the normalized root-mean-square distance from the target (legend). **C-F**. The eigenmodes corresponding to the most effectively quenched fluctuations, expected (dashed blue) and estimated by perturbations (legend as in panel B). **G**. The expected linearized dynamics matrix. **M**− **I. H**. The dynamics matrix, **M** − **I**, as estimated from the perturbation method in the 10% noise condition.

To mimic experimentally reasonable conditions, we attempt using the perturbation method by averaging over 10 repetitions of a spanning set of target deflections and allowing the network to ‘forget’ the intervention for 2 unperturbed trials. This amounts to about 1500 trials which is similar to human studies (e.g. in (9)).

With trial-by-trial fluctuations of magnitude ∼5-10% in the aforementioned distance measure around the fixed point, Fig. 1B-F show that we achieve a very close estimate of the noiseless linearized dynamics: the eigenvalue spectrum and leading eigenmodes of the linearized dynamics matrix **M** (see also the SI section “Interpreting the estimated learning dynamics in noisy network models” and Fig. S 4, also including the disentangled components of Eq. 2, the output covariance matrix **Q**, dynamics matrix **M**, and noise covariance matrix **Δ**).

When the noise fluctuations surpass 10%, the learning process by which our toy model acquires the target motion converges around mean outputs, 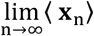, that are biased from the desired target **x**^*^ (Fig. S3 and SI section “Interpreting the estimated learning dynamics in noisy network models”). Never the less, in the 10-20% r.m.s. regime, our method extracts only slightly noisier results compared to the 5% regime above (red traces in Fig. 1B-F). The singular value spectrum, shown in Fig. 1B, has a mean error of 1.3-2.7% (SD = 1.5-3%). The greater-than-one estimated singular values can serve as a marker for the inaccuracy of the method since it indicates instability that does not exist in the learning process. Still, we get a very good approximation of the subspace near the fixed point containing the effective learning dynamics (defined by eigenmodes with singular values smaller than 1). Fig. 1C-F show the eigenmodes corresponding to the four fluctuations most effectively quenched by learning and panel H shows the full dynamics matrix. In the 5-20% r.m.s. fluctuation regime, the estimation r.m.s. errors for these eigenmodes are 1.5-2.5%, 1.6-3%, 2.9-5.3%, 4.3-7.2% (The increase in error is expected as the eigenvalues approach 1 and the effect of the learning dynamics on the fluctuations diminishes). In the case of linear gradient descent learning dynamics (Eq. 5), these eigenmodes also define the activity manifold of neurons in the network (at the output sampling rate, see methods section “Interpretation of eigenmodes and eigenvalues of the dynamics matrix M of gradient descent learning steps”). Here, the shapes of the eigenmodes in Fig. 1 reveal intrinsic properties of the rate based neural network dynamics such as oscillations indicating the periodic behavior of these models (Fig. S1).

### Example 2: a recurrent network of integrate-and-fire neurons

In the rate-based neural network we modeled additive Gaussian noise that shares the behavior time scale. An additional source of noise can arise from intrinsic stochastic properties of a spiking-neurons-based network that generates the motor drive. Spiking neurons, unlike rate-based models, may fire their action potentials with a voltage dependent probability, thus creating non-repeating single trial dynamics. Furthermore, the jitter created from probabilistic firing accumulates in a network of recurrently communicating neurons. For example, the network we describe in Methods shows a variable total number of spikes across the trials, as shown in Fig. S 5.

Comparing to the rate-based model, the simplest extension to an integrate-and-fire network is to use a readout neuron that smooths or bins the spikes of the neurons in the network to produce the output. We choose the trial n output to be **x**(t; **w**(n)) = Σ_i_**w**_i_(n) . C_i_(t), where the weights, **w**, are as before and the spike count vector of neuron at time t is 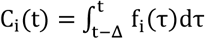 and the neuronal spike trains, 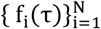, are derived from the dynamics we describe in Methods. Importantly, the inter-trial learning dynamics does not change and still follow gradient descent, Eq. 5, to converge to the desired target.

There are substantial distinctions between the rate-based network and the Integrate-and-Fire network and some important ones are discussed in SI section “Important differences between rate-based and I&F networks”. Related to testing our perturbation method we note two important distinctions. First, the noise we added to the rate-based network dynamics in example #1 had the same time binning and timescale as the desired output. The single trial dynamics in the Integrate-and-Fire case have a much finer timescale. Namely, while the duration of the network output used in the learning step (Eq. 5) is T = 0 9Sec divided into 52 time bins, it is supported by intrinsic dynamics whose fastest events, the spikes, are of ∼1 mSec. Second, unlike the rate-based model, we cannot use a ‘noise free’ network to derive the expected dynamics matrix M because the stochasticity leads to qualitatively different network dynamics and therefore to different results from Eq. 4. We therefore use an average of multiple runs, ⟨f_i_(t) ⟩, plugged into Eq. 4, as an expected result for the following analyses.

The faster timescales and intrinsically probabilistic spiking manifest in the learning dynamics matrix **M**, that operates on the coarse timescale only. Treating the network’s output as external observations of a behavioral experiment we apply our perturbation method and, in Fig. 2, show the accuracy in estimating the spectrum, leading eigenvectors, and compare the expected and estimated matrices in panels G,H. The results, shown in Fig. 2 were obtained with 100 repetitions in every dimension amounting to an order of 5000 trials. In realizable experimental paradigms that cannot reach such trial numbers it is still feasible to focus on a subset of the behavioral major variability axes (most significant dimensions of the covariance matrix **Q**).

**Fig 2.**
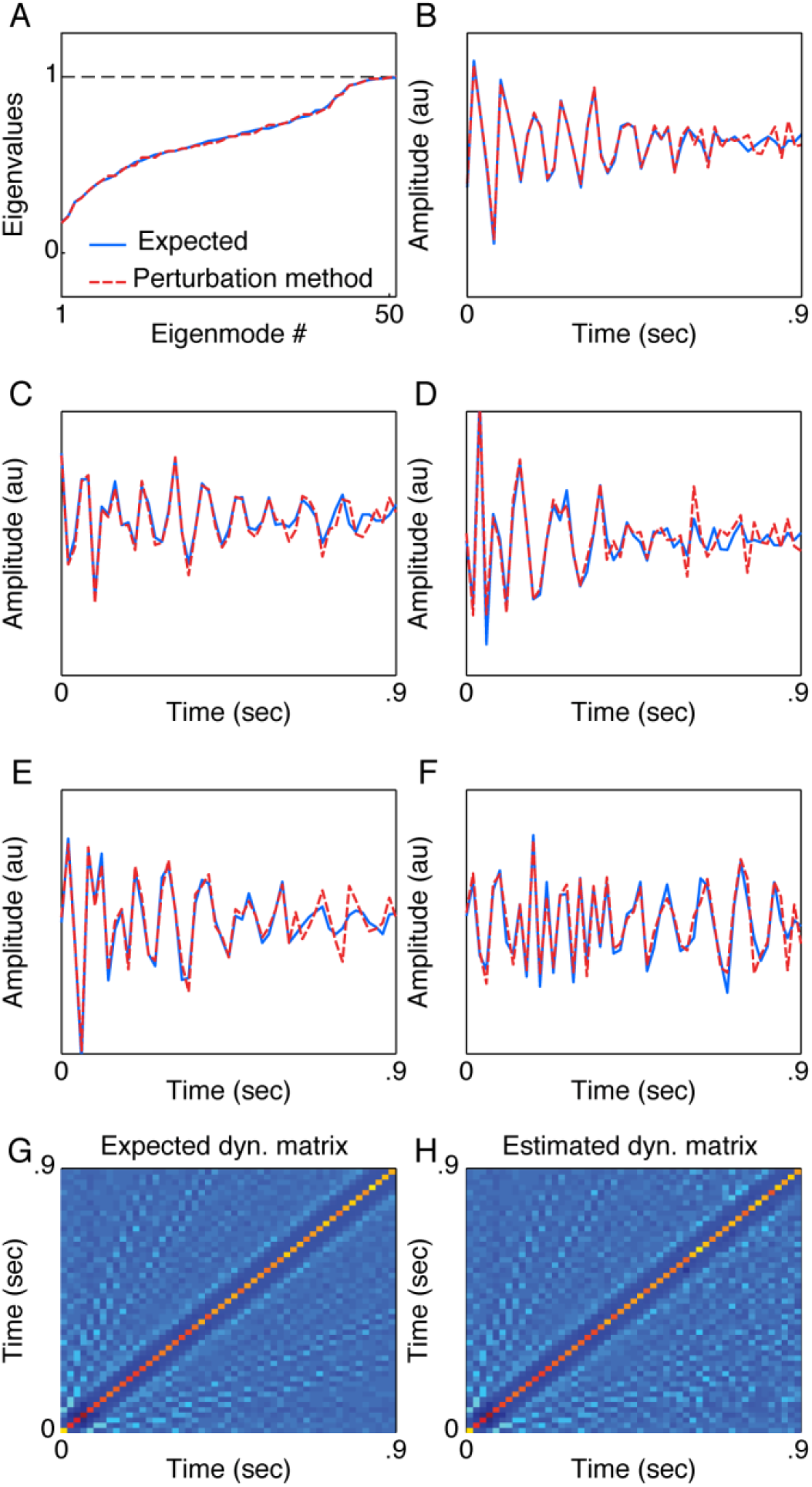
Perturbation method applied to an Integrate-and-Fire network. **A-F**. show the eigenvalue spectrum and leading eigenmodes (as described in Fig. 1) for the integrate-and-fire neural network. **G**. The expected dynamics matrix. **H**. The dynamics matrix as estimated from the perturbation method.

Comparing to the results of the rate-based model from Fig. 1, we notice the higher dimensionality of **M**, manifested in the larger number of singular values smaller than 1 in Fig. 2A. The higher dimensionality of **M** means that there are more dimensions in which the dynamics is contracting around the fixed point. In the case of linear gradient descent learning dynamics (Eq. 5), the larger dimensionality of **M** implies a higher dimensionality of intrinsic dynamics resulting from uncorrelated spike times of different neurons. This example demonstrates that faster intrinsic noise of spiking networks can be countered by harnessing more neurons to produce a reliable output target.

We also notice that the leading eigenvectors, the dimensions in which the learning is most contractive (Fig. 2B-F), have a decreasing envelope (along the within-trial time). This is an indication of the decaying level of network activity during the trial. A decaying dynamics of the generative mechanism is not trivially visible in the network’s outputs. It also implies that creating the desired output requires more dimensions, or more neurons in the network, to compensate for the unreliable late segments of the trial (Fig. S 5).

Finally, unlike the rate model network example, whose leading eigenvectors exhibited oscillatory dynamics reflecting intrinsic network properties (Fig. 1B-D), the eigenvectors of **M** in the Integrate-and-Fire example show greater complexity. These eigenvectors, shown in Fig. 2B-F, reflect neural activity fluctuations in the shorter intrinsic timescales. But, these fluctuations are close to the binning timescale and are not always fixed in frequency (c.f. Fig. 2F), suggesting aliasing of several higher frequencies caused by the binning of neural activity in the production of the output **x**(t; **w**(n)).

### Example 3: Investigating nonlinearities

So far, we demonstrated the application of our perturbation method to systems with linear learning dynamics. Perturbations are essential to unambiguous characterizations of LDS (c.f. discussion in (17)) and our framework can effectively complement these model-based parameter fitting approaches when the number of parameters is large and there is a need to explore the system’s degrees-of-freedom and prune the problem’s dimensions. However, in many common cases, such as large perturbations and within-trial responses, the learning dynamics cannot be assumed as linear.

Nonlinear learning dynamics can still converge to the neighborhood of a fixed target by reducing the execution error. Unlike the linear cases, the error gradient is a nonlinear function of the output sequence dimensions. Namely, as illustrated in Fig. 3A, a perturbation of the steady-state target, represented by the ball at the center of the error manifold, is akin to ‘kicking’ the ball. Since the learning dynamics is nonlinear, the response to perturbation will depend on the direction and magnitude of the kick. Our method can readily be used in such cases to investigate the error manifold. By changing the sets of perturbation directions we can identify purely contracting dimensions, as in Fig. 3A (i), in which perturbations are quenched along their axis with a magnitude-dependent strength, and perturbation dimensions that result in rotation dynamics, as in Fig. 3A (ii),(iii). Importantly, applying the LDS parameter estimation in this case would have yielded contradicting results if the perturbations are along direction (i) and its orthonormal complement or along directions (ii) and (iii).

**Fig 3.**
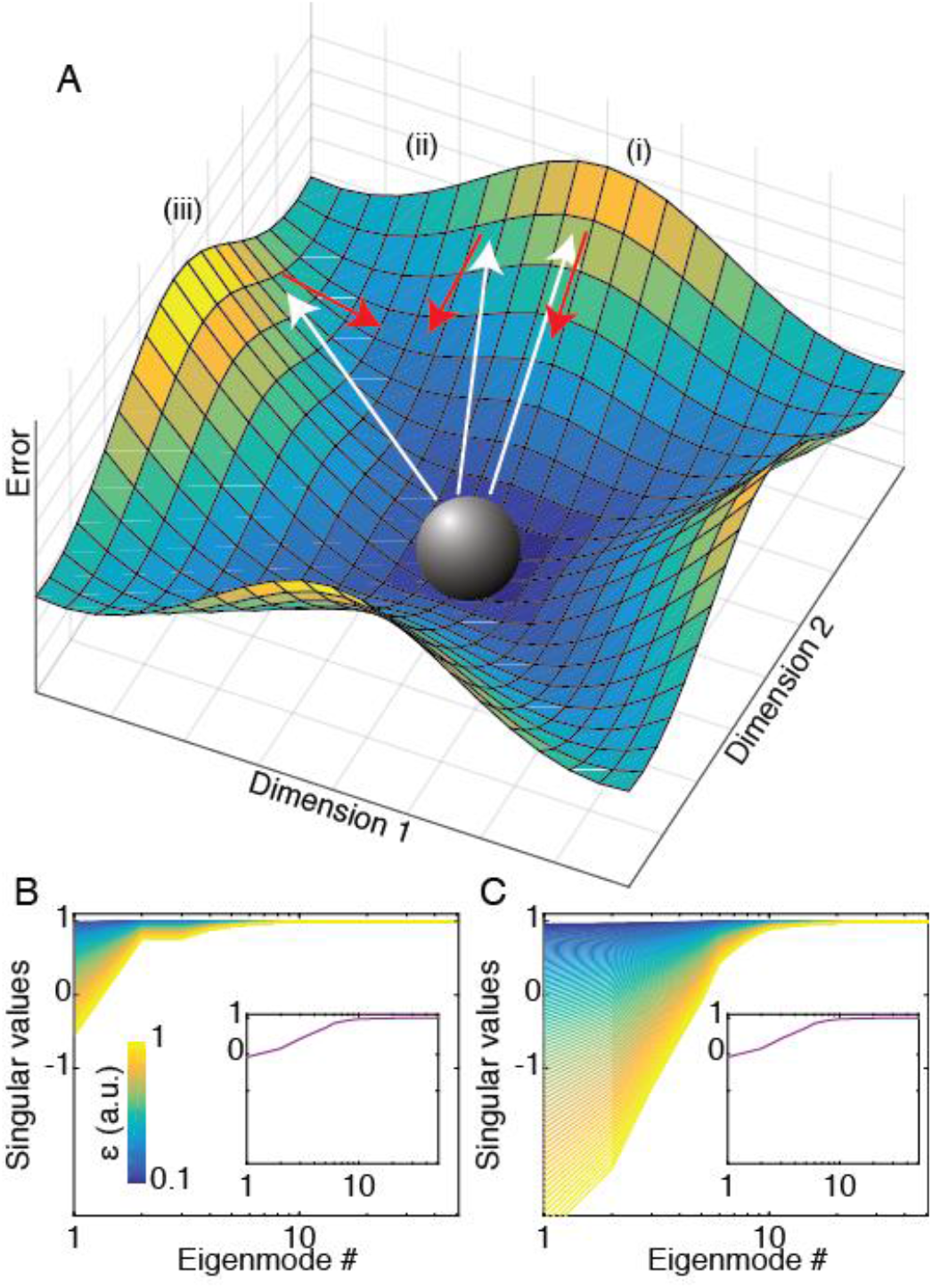
Perturbation method applied to nonlinear learning dynamics. **A**. Conceptual illustration of perturbation in a nonlinear system. Perturbations are akin to kicking the ball, which represents the fixed point in steady state at the origin of the error manifold, along the white arrows. The learning response, represented by the red arrows, depends on both the direction and magnitude of the perturbation and can be, like the linear case, simply contracting (i) or also rotating (ii,iii). **B**,**C**. The eigenvalue spectra of the linearized dynamics matrix, ***M***, extracted by perturbing a nonlinear system - the network from example #1 and a gradient learning rule on the error’s fourth power. The color bar indicates the perturbation magnitude (*ϵ*) and the insets shows the spectrum for the quadratic error case in example #1. **B**. Using the principal components of the covariance matrix, ***Q***, as perturbations. **C**. Using the standard orthonormal basis of ℝ^*D*^ as perturbations.

As an example, we repeated the simulation of the rate-based neural network but changed the learning dynamics to a gradient descent on the error’s fourth power. We applied our perturbation approach using a range of perturbation strengths (described here in arbitrary units in the range ϵ ∈ [0 1,1]) and two linearly spanning sets of perturbations. With the quadratic error function from example #1, shown in insets of Fig. 3B,C, the results of the experiments do not depend on the choice of basis and perturbation magnitude. But, in this example, the higher polynomial degree of the error function exhibits a flatter neighborhood of the target. This means that very small perturbations are under-corrected and large perturbations are over corrected. Fig. 3B,C, show the eigenvalue spectra of the linearized dynamics, extracted with the perturbation method. Indeed, with small perturbations the eigenvalues are nearly all close to 1 and the dynamics show no error correction. As the perturbation magnitude increases, the number of dimensions along which there is significant correction increases. Additionally, the strength of response to perturbations depends on the choice of basis. In Fig. 3B we used perturbations along the principal directions of the system’s covariance matrix, **Q**. But, in contrast, in Fig. 3C we used perturbations along the standard orthonormal basis. Since using this basis introduces δ -function-like errors that are very different from the neural network’s intrinsic dynamics, the response to these errors expose the nonlinearity even in weaker perturbation magnitudes. The < −1 singular values in Fig. 3C do not necessarily represent unstable dynamics but rather an over estimation of the correction that results from applying a linear response analysis to a nonlinear system.

## Discussion

Motor learning for precise execution often takes the form of trial-by-trial practicing. Investigating early steps of this acquisition process is challenging because these initial trials involve coarse corrections and non-repeating conditions. Landmark experiments used sensory-motor perturbations after the learning process converged and described behavior adaptation as linear transformations in a state space of motor primitives. This linear dynamical systems (LDS) approach is challenged when perturbations are too large or too frequent to assume linear responses and when the number of the dynamical system’s degrees of freedom is too large for reliable parameter estimation.

Motivated by experimentally realizable approaches, we proposed a method for extracting the linearized dynamics by averaging the response to small and sparse perturbations of the target motion (Eq. 3). Our approach overcomes the requirements of linearity and small number of parameters in the LDS framework. This approach is based on the assumption that the learning mechanism continues to work in steady state next to the target to prevent behavioral drift and increased variability. The perturbation method is expected to work whenever the linearization of small perturbations around the learning target is applicable.

Using examples of both noisy rate-based and stochastic Integrate-and-Fire neural networks we demonstrated that our approach is well suited for extracting the noiseless linearized learning dynamics and disentangling it from the noise covariance (in Lyapunov’s equation, Eq. 2). By estimating the linearized dynamics’ singular value spectrum and leading eigenmodes, corresponding to the most effectively quenched perturbations, we show that our approach is able to uncover key distinctions between the ‘black-boxes’ that support the within-trial dynamics, as well as the between-trial learning steps. These differences manifest in the dimensionality of the dynamics around the fixed point, determined by the singular values’ spectrum, and in the shape of the leading eigenmodes that, in the case of linear gradient descent learning, reflect intrinsic network dynamics, oscillations, and neural activity decay times. As the number of musculoskeletal degrees of freedom is much smaller than the dimension of neural dynamics, our method may tease apart the task relevant and irrelevant dimensions in stochastic optimal feedback control theories (3).

Our ‘black-box’ approach is experimentally appealing because it is both tractable and, importantly, because it is not based on specific assumptions about the learning mechanism. Strong assumptions about internal learning mechanisms can lead to designing experiments in which the behavior is highly constrained and, in many cases, unnatural and overly simple so it can be fitted by the behavior models. Our method allows taking the opposite approach because it only benefits from complex, high-dimensional, and more ethological behavior. Specifically, our method will be most efficiently utilized to study behaviors that are naturally learned to a high degree of precision like speech, birdsong, and handwriting. Such an approach may also be suitable for testing assumptions about conceptually different neural systems such as near chaos networks’ temporal stimulus response (27).

Studying the linearized component of the learning dynamics is not expected to provide a full description of internal learning mechanisms. However, the key elements and observations, obtained by our easy-to-implement method can form a basis for extrapolating into more complex, noisier, and nonlinear mechanisms. Here we have shown how this method can be used for investigating the nonlinearities in the vicinity of the learning target by exploring perturbation magnitudes and different spanning sets of directions. Understanding the learning dynamics around the behavioral fixed point will allow designing better experimental tasks and guiding in-vivo neuroscience paradigms that probe into the internal neural representations of the learning mechanisms.

Finally, beyond motor learning, we expect that our perturbation method can be used to probe other types of fixed points in complex systems. For example, we can envision self-organizing group or social structure such as social media, transportation, or homeostatic rules in gene regulation and immunology, whose target function can be perturbed to apply our approach.

## Methods

### Steady state stochastic dynamics in the vicinity of a fixed point

The variable **y**_n_ = **x**_n_ − **x**^*^ measures the deviation from the fixed point at X^*^ at the n-th trial of the learning dynamics. The stochastic linearized dynamics in the vicinity of the fixed point is described by the discrete Langevin equation (28) **y**_n+1_ = **M y**_n_ + **ξ**_n_, with i.i.d. Gaussian noise **ξ**_n_ ∼ 𝒩 (0, **Δ**). When averaged over the noise, ⟨ **y**_n_ ⟩ = 0 with covariance matrix 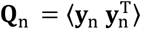.

Under the linearized dynamics, the evolution of this covariance matrix is given by 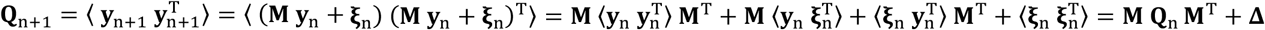, where the D X D matrix 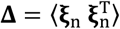 iis the noise covariance matrix. The steady-state assumption **Q**_n+1_ = **Q**_n_ = **Q** leads to the discrete Lyapunov equation (25),

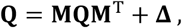

whose formal solution is **Q** = Σ_k_ **M**^k^ **Δ** (**M**^T^)^k^). In a geometric interpretation of the Lypaunov equation, the steady state covariance **Q** represents an ellipsoid volume in ℝ ^D^, which is being contracted and rotated by the dynamics matrix **M**; these changes are compensated by the volume added by the noise covariance **Δ** Readers familiar with Kalman filtering (29,30) will note that the Lyapunov equation is its steady state version.

### Extracting the dynamics matrix M by averaging over small and sparse perturbations

A target perturbation **x**^*^ → **x**^*^ + ϵ **v** applied at the end of the n-th trial changes the error from **y**_n_ = **x**_n_ − **x**^*^ to 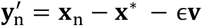, where primed variables signal the presence of a perturbation. This error changes in the next trial according to 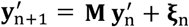, with 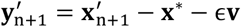. Then 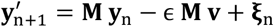, and 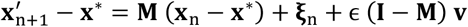.

The target perturbation is then repeated many times in a fixed direction **v**, while letting the system relax to its unperturbed target **x**^*^ for several learning steps between perturbations. Averaging across the repetitions at fixed **v** implies ⟨ **M** (**x**_n_ − **x**^*^) + **ξ** _n_)_repetitions_ → 0, leading to ⟨ **x′** − **x**^*^ ⟩_repetitions_ = ϵ (**I** − **M**) **v**.

### Linear dynamics matrix for a readout neuron trained by gradient descent

A linear neuron that outputs 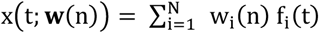 is trained to produce a desired output **x**^*^(t),0 ≤ t ≤ K . The learning algorithm is gradient descent on the squared error 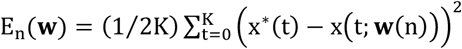. Gradient descent 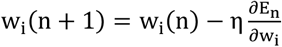 with a learning rate η implies

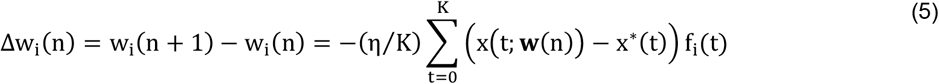

It is interesting to note that that the stability of the learning fixed point at **x**^*^(t), which follows from the Hessian matrix, 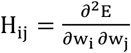, is determined by the dynamics of the recurrent network, since 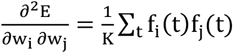. The rank of the Hessian is limited by the discrete duration K of each trial.

Assume that the learning process converged to **w**(n) = **w**^*^; we then perturb the target function to **x**^*^(t) = **x**^*^(t) + ϵV(t) The next learning step results in 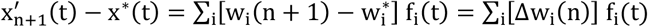. Substituting **Δ** w_i_(n) = −(η/ K) Σ_τ_ (**x**(τ**w**^*^(n)) − **x**^*^(τ) − ϵV(τ)) f_i_(τ) as per Eq. 5 results in

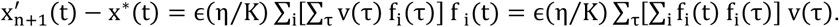

In the noiseless case considered here, this leads to the identification of the linearized dynamics matrix of Eq. 4 because by Eq. 3 it equals ϵ (**I** − **M**)**v**.

### Interpretation of eigenmodes and eigenvalues of the dynamics matrix M of gradient descent learning steps

Eq. 4 defines the dynamics matrix **M** for the gradient descent algorithm using the squared error as in Eq. 5 above. If **M** is symmetric (e.g. in Fig. 1) its eigenmodes, 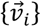 satisfying 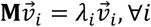, are an orthonormal spanning set of ℝ^D^ (the motor output space). For a noiseless neural network with N neurons producing the output traces 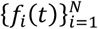, we can span the output of each node by substituting 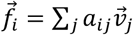 where a_*ij*_ are real valued coefficients. Plugging into Eq. 4 we get 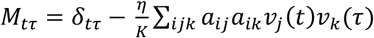 . To satisfy the eigenmode properties we therefore demand 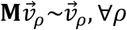 leading to 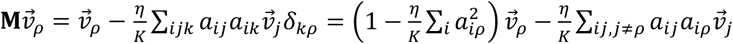. Since the last term in this expression is orthogonal to 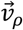 it must be zero to allow the right-hand side to be a scalar multiplication of 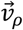. Additionally, the first term, 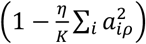, is always smaller or equal to 1. Thus, if 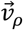 has the eigenvalue 1 we must have 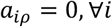.

The interpretation of this algebra is that the neural activity, 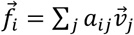, can be spanned by a subset of eigenmodes whose eigenvalues are smaller than 1. Specifically,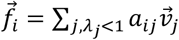.

In other words, the eigenmodes of **M** with eigenvalues smaller than 1 (and larger than −1) define the manifold of the neural activity if the learning step is gradient descent on the squared error (Eq. 5).

### Neural network simulation: firing rate neurons

Consider a recurrent neural network with full connectivity. The firing rate of each neuron 1 ≤ i≤ N is f_i_(t) = tan h(r_i_(t)), with equations of motion

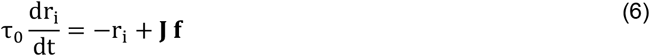

The elements of the N × N connectivity matrix **J** are chosen randomly,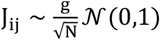. Here we use N = 1000 neurons and g = 1 2. For these choices, the dynamics of the recurrent network is between simple periodic and chaotic ((31), also illustrated in Fig. S 1). In this regime the linear readout neuron can be trained to produce a wide range of outputs (23).

Neurons in the recurrent network are initialized with 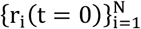 independently drawn from the Gaussian distribution 𝒩 (0,1/2). After initialization, the network runs for a 10 seconds ‘burnout’ time. Then, learning trials are simulated using snippets of duration T = 2s during which the recurrent network evolves in time and provides inputs to the readout neuron. At a temporal resolution of δt = 40ms, a snippet corresponds to K = T/ δt = 50 discrete time steps following the trial onset. An interval of T = 2s during which the network continues to follow its intrinsic dynamics is allowed to elapse before the following snippet to simulate the inter-trial interval (ITI).

The beginning of each snippet corresponding to a trial is indicated by a neural ’go-signal’, implemented here through the addition of a constant activation r_0_ = 10 to all 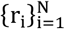 at the beginning of each trial. This pulse of increased activation is the same for all neurons and across all trials, and its application at trial onset primes each trial snippet with a degree of commonality in initial conditions. Fig. S 1 shows that it takes ∼0.3 sec in simulation time for the network to return to the region of its steady state dynamics after the pulse onset.

To evaluate the effect of different initial conditions separately from the learning procedure, we repeat the network initialization and ‘burnout’ steps, described above, 25 times. We use the first snippet of T = 2s after the burnout to compute the **M** matrix, its eigenvalues and eigenvectors, to establish their stability (Fig. S 2). In the SI section entitled “Initial conditions, ‘go’ signal, and steady state dynamics of the rate-based network simulations” we discuss the behavior of the simulated network at trial initiation and at asymptotic times.

### Learning the weights of the readout neuron in the presence of noise

We chose K = 50 time steps for the duration of a trial, and a learning rate η = 0.02 for the gradient descent algorithm of Eq. 5. We incorporated additive noise to the equations of motion of the neural networks, Eq. 6,

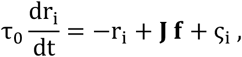

where the 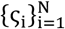 were randomly and independently chosen from a Gaussian distribution 𝒩 (0, σ^2^), with standard deviation σ = 0 0,0 002,0 00s,0 01 This range was chosen because the added noise biases the learning process from the desired target motion. In the SI section “Interpreting the estimated learning dynamics in noisy network models” we detail this process and show that when a = 0 01 the learning process converges around a mean output biased from the desired target motion by about 25% (defined as the vector norm of the difference divided by the vector norm of the target, Fig. S 3C). The perturbation method we propose in this manuscript performs less accurately at this high noise condition, so we did not explore larger noise (σ > 0 01).

The simulations of networks of firing rate neurons were done with MATLAB (Mathworks).

For each value of σ, we used 1000 consecutive learning trials to estimate the output covariance around the target, *Q*_*tτ*_= ⟨*y*_*n*_(*t*)*y*_*n*_(*τ*) ⟩. Then, we calculated the eigenvectors and eigenvalues of **Q**. The eigenvalues were used to estimate the dimensionality of the output manifold (Fig. S 3B) and the eigenvectors were used as perturbation directions for the results in Fig. 1 and Fig. 3.

For each of the noise conditions we calculate the norm of the mean deviance from the target, ∥ ⟨*y*_*n*_(*t*) ⟩∥, and the mean of the norm, ⟨∥ *y*_*n*_(*t*) ∥ ⟩. Fig. S 3C,D show those values normalized by the norm of the target, ∥ **x**^*^ ∥.

To mimic experimentally plausible conditions, we apply the perturbations every 3^rd^ trial and repeat every perturbation direction 10 times.

### Neural network simulation: integrate-and-fire neurons

Unlike the rate-based neuron model (Eq. 6), the Integrate-and-Fire (I&F) neuron model transmits millisecond-scale spike times to its postsynaptic targets. Here we constructed a network of integrate-and-fire neurons with output timescales comparable to those observed in the simulations of networks of rate-based neurons. Qualitative differences between the rate-based and I&F networks are further discussed in the SI section entitled “Important differences between rate-based and I&F networks”.

The simulations of networks of integrate-and-fire neurons were implemented in C and parallelized with cluster computing (LSF).

The dynamic variables that characterized the state of each neuron 1 ≤ i ≤ N are listed in Table 1. The parameters used in the network simulation are listed in Table 2.

**Table 1.** State variables of integrate-and-fire neurons.

| variable | symbol | units |
| --- | --- | --- |
| Membrane potential of neuron i | $V_i$ | mV |
| Conductivity coefficient of synapses from neuron j | $P_j$ | dimensionless |

**Table 2.**
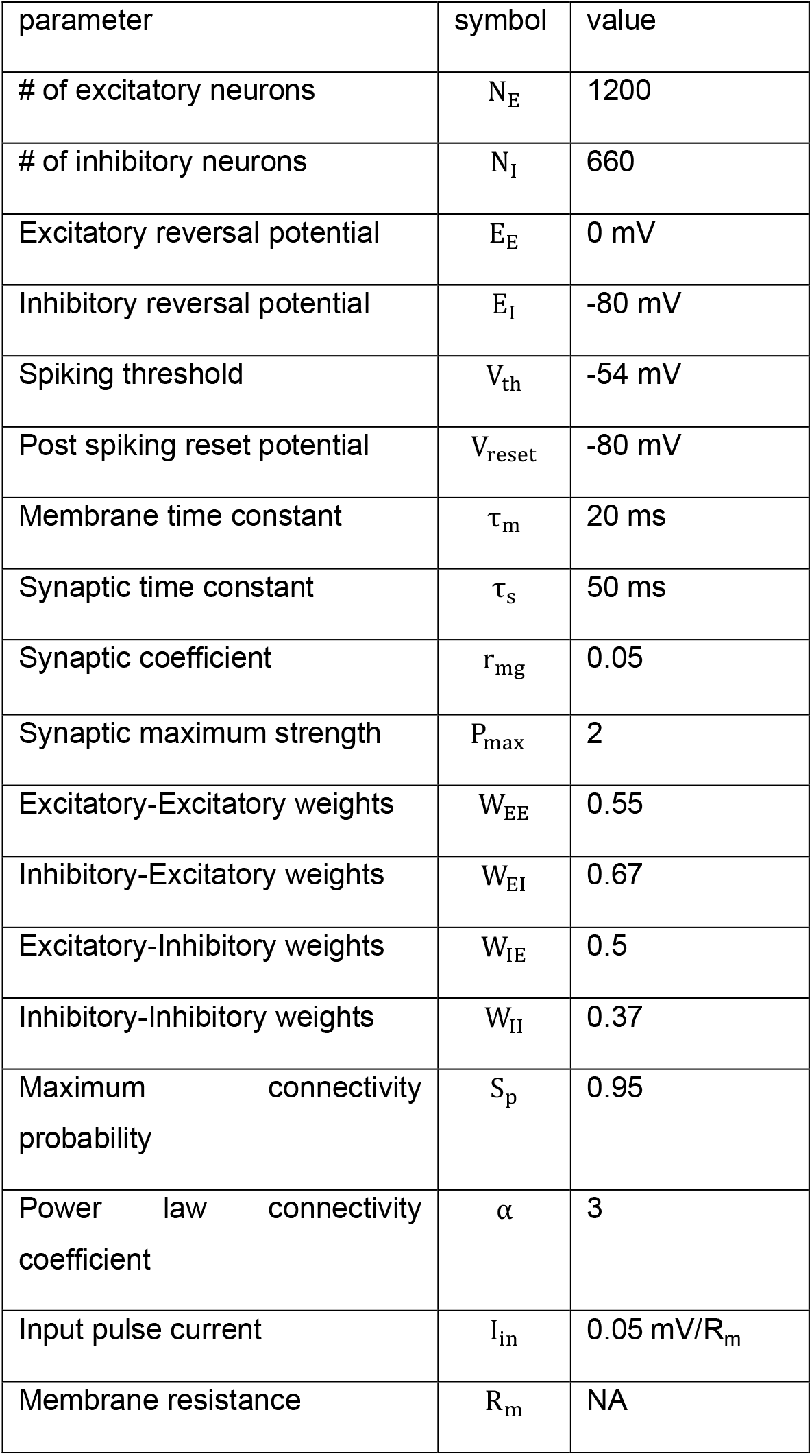
Model parameters for network of integrate-and-fire neurons.

#### Network connectivity

The connection J _ij_ from neuron j to neuron is chosen as follows. There are no self-connections:J _ii_ = 0 for all 1 ≤ i ≤ N. For i≠ j, the existence of a connection is stochastically set according to a power law distribution with coefficient α = 3. The strength of existing connections depends on whether the neurons belong to the same class: E (excitatory) or I (inhibitory). We use indices a, b ∈ {E, I} to formulate the biased rule 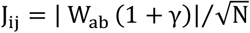; here γ is a stochastic variable drawn from 𝒩 (0,1) Note that all connections are positive; the excitatory or inhibitory nature of a connection is determined by the corresponding reverse potential, controlled by the presynaptic neuron (see the equation for the update of V_i_(t) below).

#### Equations of motion

The membrane potential of all neurons 1 ≤ I ≤ N is updated simultaneously at time intervals dt according to

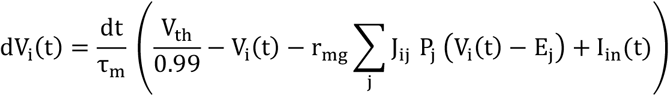

Neuron i will emit a spike in time bin [t − dt, t] if tanh 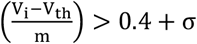. We used m = 0.25 mV and σ ∼ 𝒩 (0,0.0001).

After emitting a spike, the membrane potential and the and synaptic conductivity of neuron are reset to

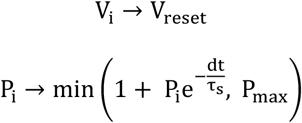

When no spikes are emitted, synaptic conductivities decay. If their value falls below 10^−5^, they are set to 0:

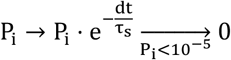

## Acknowledgments

The authors thank Alon Rubin, Ori Maoz, Nathan Perkins, and Misha Tsodyks for valuable suggestions and comments.

## Supporting Information (SI)

### Initial conditions, ‘go’ signal, and steady state dynamics of the rate-based network simulations

In this manuscript we focused on rate-based network simulations both because rate-based frameworks are leading approaches in in-vivo and in-silico studies and because of the established theoretical understanding of these models. As substrates of motor execution, neural networks have two necessary properties; (1) The ability to support diverse outputs, and (2) the ability to support stable outputs.

For the network models we consider, these properties depend on the coupling between neurons, on the initial conditions, and on the noise added to the equations of motion (Eq. 6).

#### Coupling strength has opposing effects on output diversity and stability

We chose rate-based networks of N = 1000 neurons with coupling coefficients sampled from a Gaussian distribution,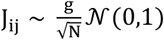. The coupling constant in such models, *g*, determines their steady-state dynamics and response to transient inputs (31). These properties can be visualized using time embedding (Fig. S 1A). This representation reveals that below *g* = 1 the network is quiescent, between *g* = 1 and *g* = 1 2 there is a range of periodic behavior, and above *g* = 1 2 the network approaches chaotic behavior. (the range of coupling strength with periodic dynamics will narrow as N increases and disappear as N → ∞)

In response to an activation input (‘go’ signal) of *r*_0_ to all the nodes (see methods) the models will display a decaying transient to their steady-state manifold (Fig. S 1B). The pulse increases the similarity between model trajectories with different initial conditions, but this similarity will quickly diverge in networks close to or in the chaotic regime. Thus, larger coupling increases the networks’ ability to span diverse outputs but decreases its reliability due to increased sensitivity to initial conditions.

**Fig S1.**
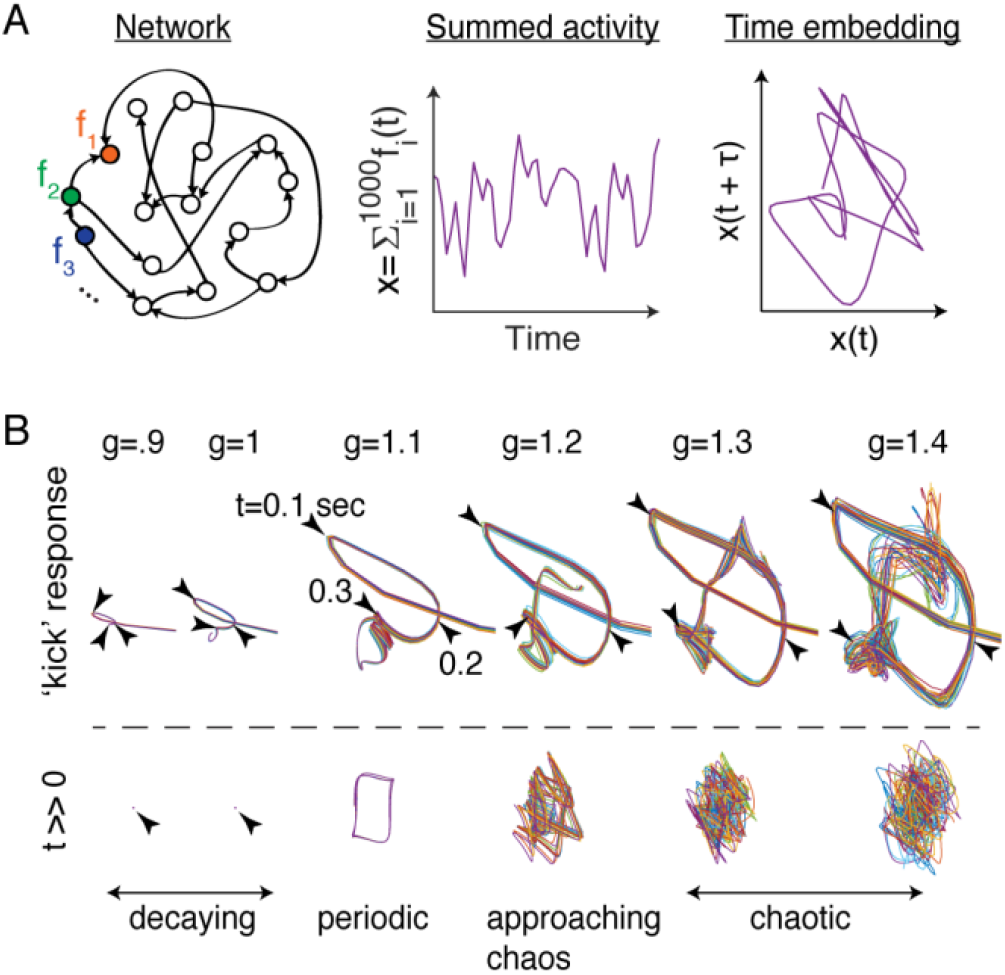
Time embedding of rate-based network activity reveals its steady state dynamics and response to transient impulse inputs. **A**. The output of each node in the network, 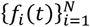, is summed to create a representative activation variable, **x**(t), equivalent to the mean firing rate. The time evolution of *x*(*t*), is embedded in 2d by plotting all points (*x*(*t*), *x*(*t* + τ)) for all *t* and a given τ (80mSec). **B**. The connectivity matrix used to create Fig. 1, was uniformly rescaled to different values of the coupling constant *g*. The network was initialized in 20 random initial conditions (colors, see methods) and simulated for 10 seconds ‘burnout’. Then, a ‘go’ activation of *r*_0_ = 10 was added to all nodes in the network. The first 1 second of the networks’ transient response is shown above the dashed line using 2d embedding. Markers show the 0.1,0.2, and 0.3 sec points. The bottom shows another 1 second of the embedded trajectory at large time (t=10 seconds). Markers show a point at the origin of a quiescent network when g ≤1.

#### A ‘go’ signal is necessary for stable learning dynamics

We chose *g* = 1 2 in this work because the network’s dynamics is not simple periodic and a single readout neuron that outputs a scalar linear combination of the activity of all neurons can learn diverse outputs and approximate the target motion we imposed (in Fig. 1, similar to (23)). In this setting the initial conditions significantly affect the firing rate dynamics of the neurons, 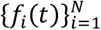, in the time snippet of duration T used to drive the motor output 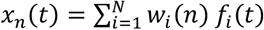 in the n^th^ trial Eq. 4 allows calculating the expected linearized <u>trial-to-trial</u> learning dynamics matrix, **M**, from the network <u>within-trial</u> dynamics 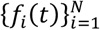. If the network dynamics is sensitive to initial conditions, we can expect instability in **M** in different trials, demonstrated through the instability of its eigenvalues and eigenvectors. This sensitivity is shown in Fig. S 2B,E. The effect of variability in initial conditions can be mitigated by adding an activation pulse (or ‘go’ signal). Fig. S 2A,D show the spectrum and leading 3 eigenmodes of **M** calculated from the dynamics of the network we used in Fig. 1 with random initial conditions and an added activation pulse – demonstrating that the reliable network outputs also lead to reliable learning dynamics. Fig. S 2C,F show the effect of increasing the coupling coefficient **g**. In the chaotic regime the activation ‘go’ signal is not enough to create reliable outputs because the network dynamics is too sensitive to its initial conditions. This instability leads to unstable learning dynamics. Specifically, certain eigenvalues become smaller than −1 which means diverging trial-to-trial dynamics.

**Fig S2.**
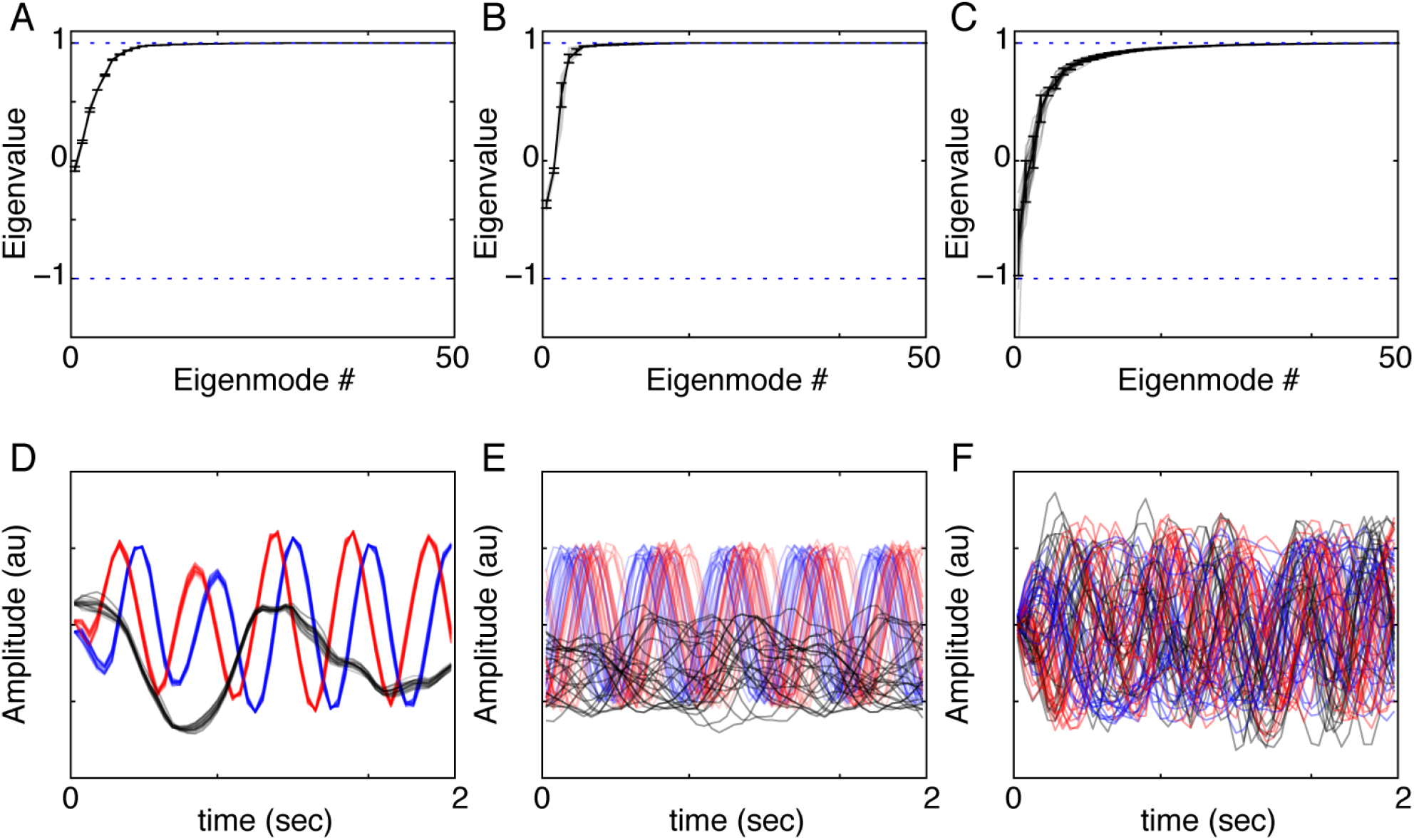
The reliability of the learning matrix M computed from the activity of a recurrent rate based neural network depends on its initial conditions and connectivity strength. To evaluate the effect of different initial conditions separately from the learning procedure, a noiseless rate-based network with the same connectivity matrix as in Fig. 1 is initialized 25 times with different i.i.d initial conditions 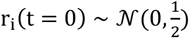. The rate model runs for 10 seconds simulated ‘burnout’ time. Then, an input pulse is added to all neurons and the network’s output in the following 2 seconds is used for calculating the learning dynamics matrix, **M** (using Eq. 4). The matrix’s eigenvalue spectrum and 3 most learning-affected eigenvectors (ordered − blue, red, and black) are compared in the following conditions: **A**. Identical conditions to Fig. 1 Gray curves show the eigenvalue spectrum of the matrix **M**. The black line and error bars are the mean and standard deviation of the gray curves. **B**. Same as A but with no added pulse. **C**. Same as A but with half the pulse size and the connectivity matrix uniformly rescaled to have the mean coupling constant **g** = 1.5 and not 1.2 as in A,B **(D-F)** The 3 most learning-affected eigenvectors corresponding to A-C respectively.

### Interpreting the estimated learning dynamics in noisy network models

We modeled noise by adding a random component to the networks’ equation of motion (Eq. 6) drawn i.i.d from a Gaussian distribution. The noise term is integrated during the networks’ temporal evolution and impacts the output of the readout neuron (Fig. S 3A). In larger noise conditions, the trial-by-trial fluctuating outputs, ***x***_*n*_(*t*), occupy higher dimensional volumes after the learning process converges - exceeding the manifold in which the learning dynamics is effectively contracting (Fig. S 3B). In these conditions, the noise can both cause larger output variability (Fig. S 3D) and, importantly, a bias of the mean output from the desired target motion, **x**^*^ (Fig. S 3C).

**Fig S3.**
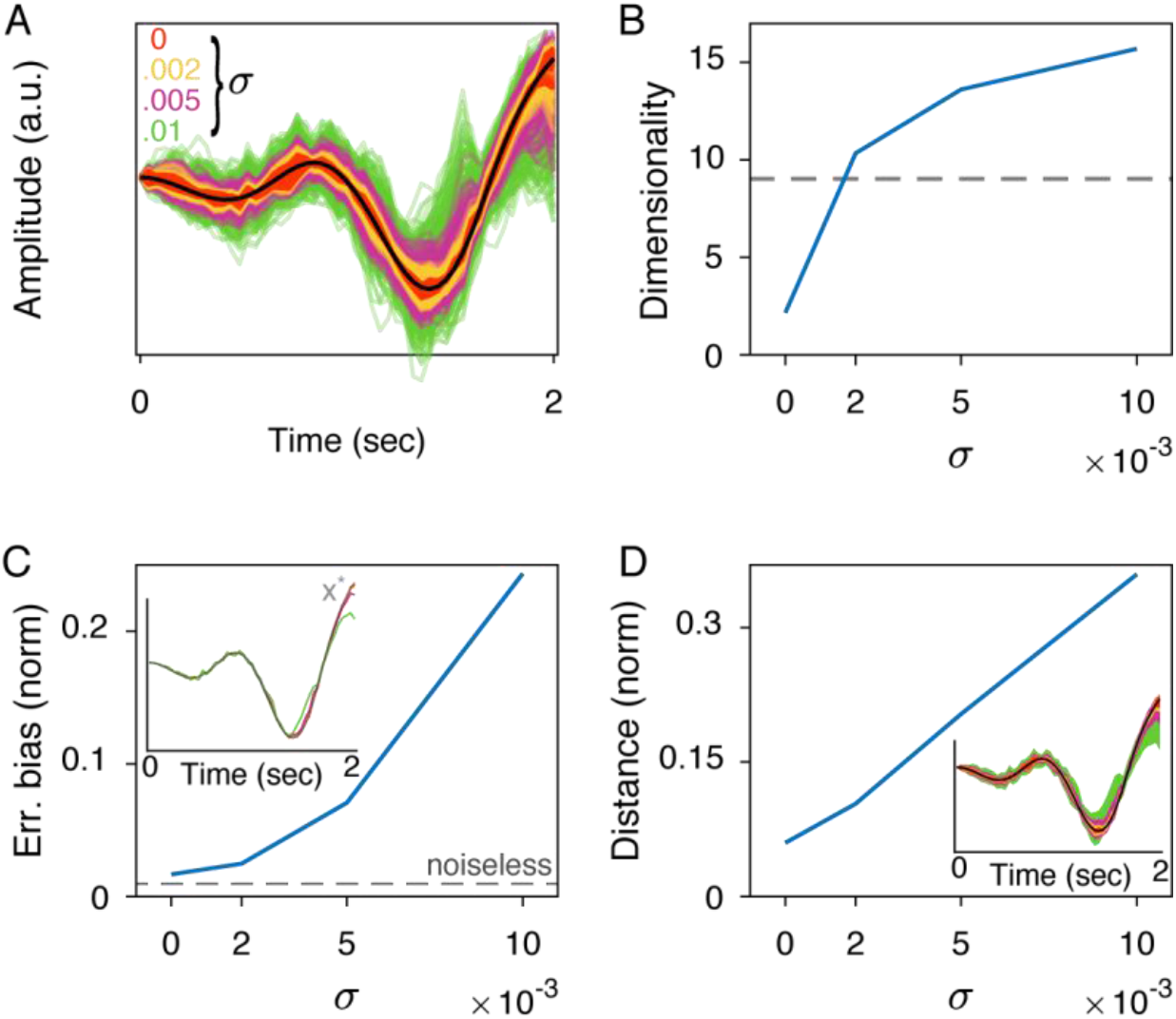
Adding Gaussian noise to the rate-based equations of motion increases the outputs’ steady-state dimensionality, bias, and variability. **A**. After the learning converges to the vicinity of the target motion (**x**^*^, black line), the variability of rate-based model outputs depend on the standard deviation of the Gaussian noise added to the equation of motion (σ, color coded. x-axis in panels b-d). **B**. The dimensionality of outputs, **x** ∈ ℝ^*D*^, is estimated by interpolating the number of covariance matrix eigenvalues required to explain 95% of the variance. Dashed line shows the number of eigenvalues smaller than 0.95 of the noiseless dynamics matrix, **M. C**. The normalized average output error (y-axis, ∥ ⟨**y**⟩∥ / ∥**x**^*^∥, where **y** = **x** − **x**^*^) shows that, in this example, the noise biases the outputs. Dashed line shows the minimal error achieved with a noiseless network with fixed initial conditions. Inset, the average outputs used to calculate the errors. (colors matching panel A. Gray marks the true target motion **x**^*^). **D**. The normalized mean distance (y-axis, ⟨∥ **y**∥ ⟩/ ∥**x**^*^∥) shows that the noise increase errors almost linearly with σ. Inset, shaded areas show the 1 STD outputs used in calculating the error r.m.s. (colors matching panel A).

The bias in the network mean output from the target motion limits the conditions in which Eq. 4 can be used to predict the expected linearized learning dynamics matrix **M**. (because the averaging across trials deviates from the firing rate dynamics of the neurons, 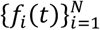, of a noiseless network initialized at r_i_ = r_0_, ∀*i* ∈ {1,... N}). In our noisy rate-based neural networks, using the perturbation closely estimated **M**’s spectrum and leading eigenmodes even when the mean network output deviated from the desired target by 7% and fluctuated in typical distances equivalent to 20% of the target’s norm (as defined in the main text for σ = 0 00s in Fig. 1B-F, Fig. S 3C,D, Fig. S 4A,B). In smaller noise conditions, e.g. σ = 0 002 or 10% fluctuations, the perturbation method estimates the matrix **M** with high precision and, using Lypunov’s equation (Eq. 2) and the measurable output covariance matrix **Q** (Fig. S 4C), allows extracting the noise covariance matrix **Δ** (Fig. S 4D).

We note that noise can also impact the trial-by-trial learning dynamics. Unlike noise in the network’s equations of motion, affecting the <u>within-trial</u> dynamics, adding a noise term to the learning step will affect the <u>between-trials</u> weight adaptation. For example, a noise term, ζ_i_, can be added to the weight update (Eq. 5),

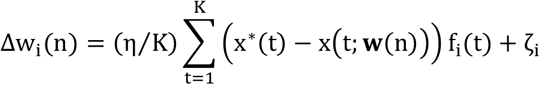

However, if the added noise, ζ_i_, has zero mean then it will not bias the network outputs from the desired target and just trivially increase the errors. For that reason, and for clarity, we did not treat such noise in this manuscript.

**Fig S4.**
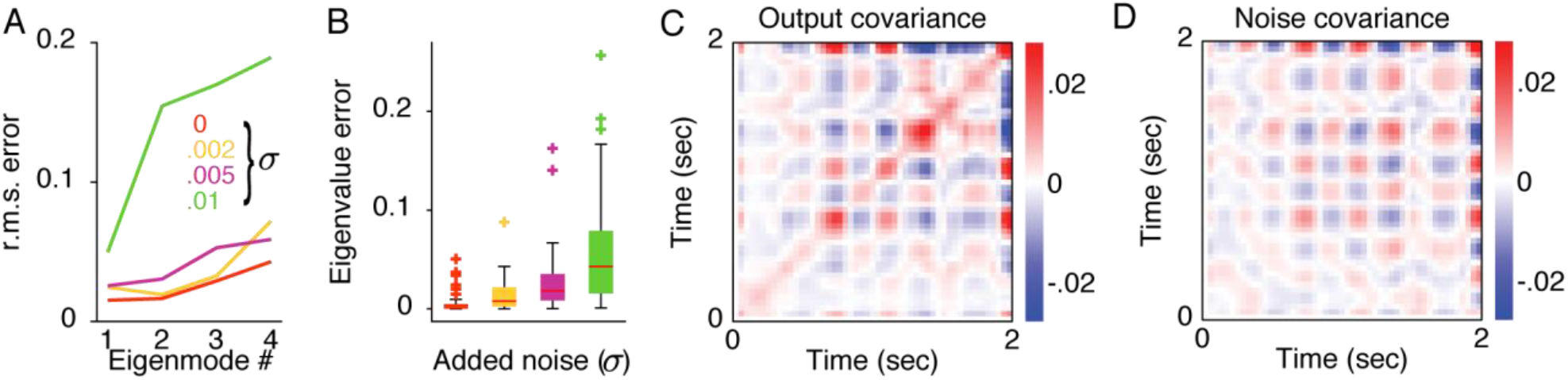
Perturbation method estimates of the dynamics matrix are used to extract the discrete noise covariance in Lyapunov’s equation. **A**. The estimation error of the 4 eigenmodes of the dynamics matrix M in Fig. 1C-F is summarized by the root-mean-square difference between the expected and estimated vectors (y-axis). Colors show different variance levels (σ, matching Fig. 1) of the noise added to the equation of motion (Eq. 6). **B**. Boxplot summarizes the eigenvalue estimation errors in Fig. 1B (y-axis, showing the difference between the expected and estimated values). Boxes show the 0.25,0.75 quantiles and whiskers show errors extending another 1.5 interquartile difference. Outlier markers show the rest of the data. **C**. The output covariance matrix **Q**, in the 10% noise example in Fig. 1H (σ = 0 002). **D**. The noise covariance matrix **Δ**, calculated from Eq. 2 using the measured **Q** and estimated **M** in Fig. 1H.

### Important differences between rate based and I&F networks

When simulating an integrate-and-fire neural network, we start with a network that contains both excitatory and inhibitory neurons. This differs from the rate model, in which the connectivity matrix, **J**, was chosen from a Gaussian distribution without limiting the sign of interaction. This means that the same unit can be both inhibitory and excitatory. Such an abstraction may apply for the rate model, where the units are not strictly regarded as neurons, but with integrate and fire models we limit ourselves to the realistic network.

Another difference is the inherent noise in the integrate and fire network that results from the influence of spike timings. In our model we introduced noise by imposing a probabilistic spiking voltage threshold. This means that each modeled neuron elicited a spike around a threshold voltage with a probability that depended on the voltage.

However, when building the integrate and fire model, we wish to maintain several similarities to the rate model:

- Response duration - We stimulate the network with an impulse current for 20mSec but demand a response that lasts 1-2 seconds and decays.
- Response dimensionality - the binned spikes vectors, 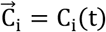, need to reliably span a high dimensional space that will allows convergence to the desired output.
- Noise properties - Inherently, the variability of the single neuron output, C_i_(t), across repeated input pulses, increases with t. Specifically, the variability in the first bins is much smaller than the last bins because of the additive effect of the jitter in spike times. The reliability, mentioned above, means that each of the single neuron responses, 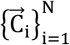, should not move too much because of the noise. Most importantly, the responses duration needs to vary in a controlled manner and always decay before the next input impulse (we keep the continuous learning simulation).
- Parameter robustness - much like the rate models, we want our simulation to be somewhat robust to parameters changes. It seems unrealistic that the slightest change in parameter values or neuron numbers should drastically change the network behavior.

As it turned out, meeting these demands is not straight forward. Using randomly connected excitatory and inhibitory neurons results in a network that has either a fast decaying response or a sustained activity response to input impulses. Fine tuning the network may result in a long decaying response but will be unreliable under noise and would most likely lack the dimensionality of the response and will of course defy the robustness requirement.

#### Integrate-and-Fire model modifications

We introduced two modifications that yielded the desired result:

1. <u>Power law connection probability</u>: We, started with the Gaussian synaptic strength distribution but decided to set a connection to zero (no connection) between a neuron to itself and also randomly with a power law distribution (pr0b(**x**) ∼**x**^−α^ on the closed set [0 01,0 9s]).
2. <u>External excitatory background activity</u>: We introduced an external input to all neurons that drove them close to the spiking threshold. 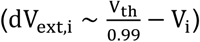

The resulting model provided the desired properties and is described in the main text.

### Variability of spike counts in an Integrate-and-Fire model

This variability, here quantified as the standard deviation of the total number of spikes in 10 mSec time bins, increases along the trial duration, t ∈ [0,1], because of the accumulated jitter and drops to zero at t→1 because the activity in the network decays.

**Fig S5.**
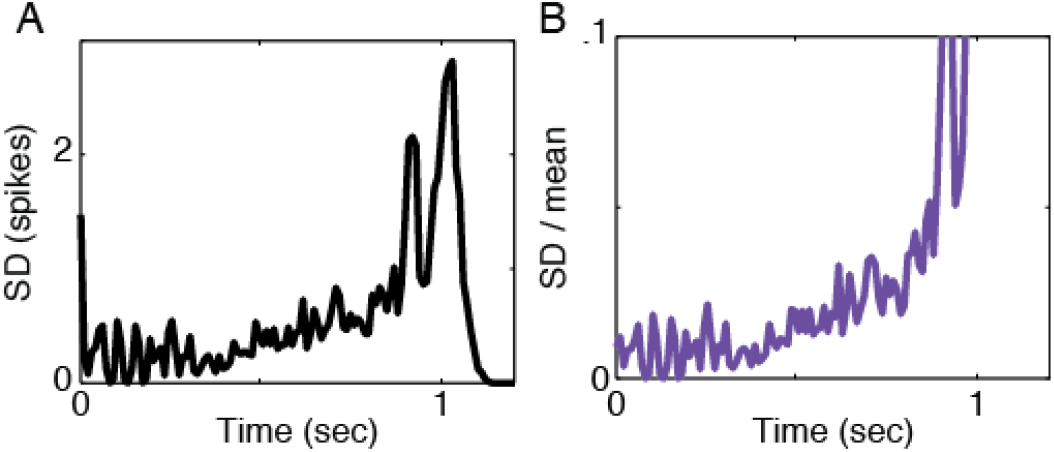
The variability of spiking neuron networks’ outputs. **A**. The standard deviation, across trials, of the number of spikes per time bin as generated by an integrate and fire network model. **B**. The ratio between the standard deviation (from A) and the mean spiking rate per bin in the same simulation.

To make sure the learning process converges, we used a segment of 0.9 Sec from each trial to produce the output of the network. As Fig. S 5A,B show, the rest of the network’s output in t ∈ [0 9,1] decays while the variability increases (SD / mean increases). This approach can be considered as using the first segment of the simulated trial for learning to produce a reliable output, swinging a golf club for example, and letting the unreliable ending of the motion to take no effect in learning because the goal has already been achieved (the golf ball was hit).

